# Chromatin Profiling of CBFA2T3-GLIS2 AMLs Identifies Key Transcription Factor Dependencies and BRG1 Inhibition as a Novel Therapeutic Strategy

**DOI:** 10.1101/2023.08.30.555598

**Authors:** Samantha Kaonis, Jenny L. Smith, Neerja Katiyar, Morgan Merrill, Tiffany Hyelkma, Stephanie Namciu, Quy Le, Ekaterina Babaeva, Takashi Ishida, Shelli M. Morris, Emily Girard, Suzanne Furuyama, Rhonda Ries, Irwin Bernstein, Soheil Meshinchi, Steven Henikoff, Michael Meers, Brandon Hadland, Jay F. Sarthy

## Abstract

Oncogenic fusions involving transcription factors are present in the majority of pediatric leukemias; however, the context-specific mechanisms they employ to drive cancer remain poorly understood. CBFA2T3-GLIS2 (C/G) fusions occur in treatment-refractory acute myeloid leukemias and are restricted to young children. To understand how the C/G fusion drives oncogenesis we applied CUT&RUN chromatin profiling to an umbilical cord blood/endothelial cell (EC) co-culture model of C/G AML that recapitulates the biology of this malignancy. We find C/G fusion binding is mediated by its zinc finger domains. Integration of fusion binding sites in C/G-transduced cells with Polycomb Repressive Complex 2 (PRC2) sites in control cord blood cells identifies *MYCN, ZFPM1, ZBTB16 and LMO2* as direct C/G targets. Transcriptomic analysis of a large pediatric AML cohort shows that these genes are upregulated in C/G patient samples. Single cell RNA-sequencing of umbilical cord blood identifies a population of megakaryocyte precursors that already express many of these genes despite lacking the fusion. By integrating CUT&RUN data with CRISPR dependency screens we identify *BRG1/SMARCA4* as a vulnerability in C/G AML. BRG1 profiling in C/G patient-derived cell lines shows that the *CBFA2T3* locus is a binding site, and treatment with clinically-available BRG1 inhibitors reduces fusion levels and downstream C/G targets including N-MYC, resulting in C/G leukemia cell death and extending survival in a murine xenograft model.

## Introduction

Large-scale cancer genomics studies have identified fusion oncoproteins as the primary genomic lesions in pediatric acute leukemias. These fusion events often join hematopoiesis-specific transcriptional co-activation domains to DNA binding domains from non-hematopoietic transcription factors. While fusion oncoproteins are a recurrent theme in pediatric leukemias, the chromatin-centric mechanisms they use to promote oncogenesis remain poorly characterized in many cases. Further, they display tremendous context specificity, often driving distinct subsets of leukemias associated with markedly different prognoses.

CBFA2T3-GLIS2 fusions are associated with childhood-restricted, treatment-refractory acute megakaryocytic leukemias^1^. CBFA2T3 (ETO2) is an ETO transcription factor family member that contains three NHR (*nervy* homology region) domains that mediate interactions with other TFs including those from ERG and RUNX families^1,2^. A fourth NHR domain contains a MYND (myeloid, *nervy* and Deaf-1) domain that binds N-CoR and SMRT co-repressors (Fig. 1a). GLIS2 is a zinc finger (Zf)-containing transcription factor primarily expressed in kidney where it promotes normal tissue homeostasis^3^. Mutations in *GLIS2* are associated with nephronophthisis and it has no established role in hematopoiesis^3^. The CBFA2T3-GLIS2 fusion arises from an inversion of chromosome 16, giving rise to a fusion transcript containing 5’ exons from *CBFA2T3* including the three NHR domains and the 3’ end of *GLIS2* containing the ZNF domains and a transactivation domain. Fusion expression is driven by the promoter and enhancers of the *CBFA2T3* locus^4^. One wildtype allele of *CBFA2T3* is also expressed in these leukemias, though a role for the wildtype protein in C/G AML leukemogenesis has yet to be fully elucidated.

**Figure 1.**
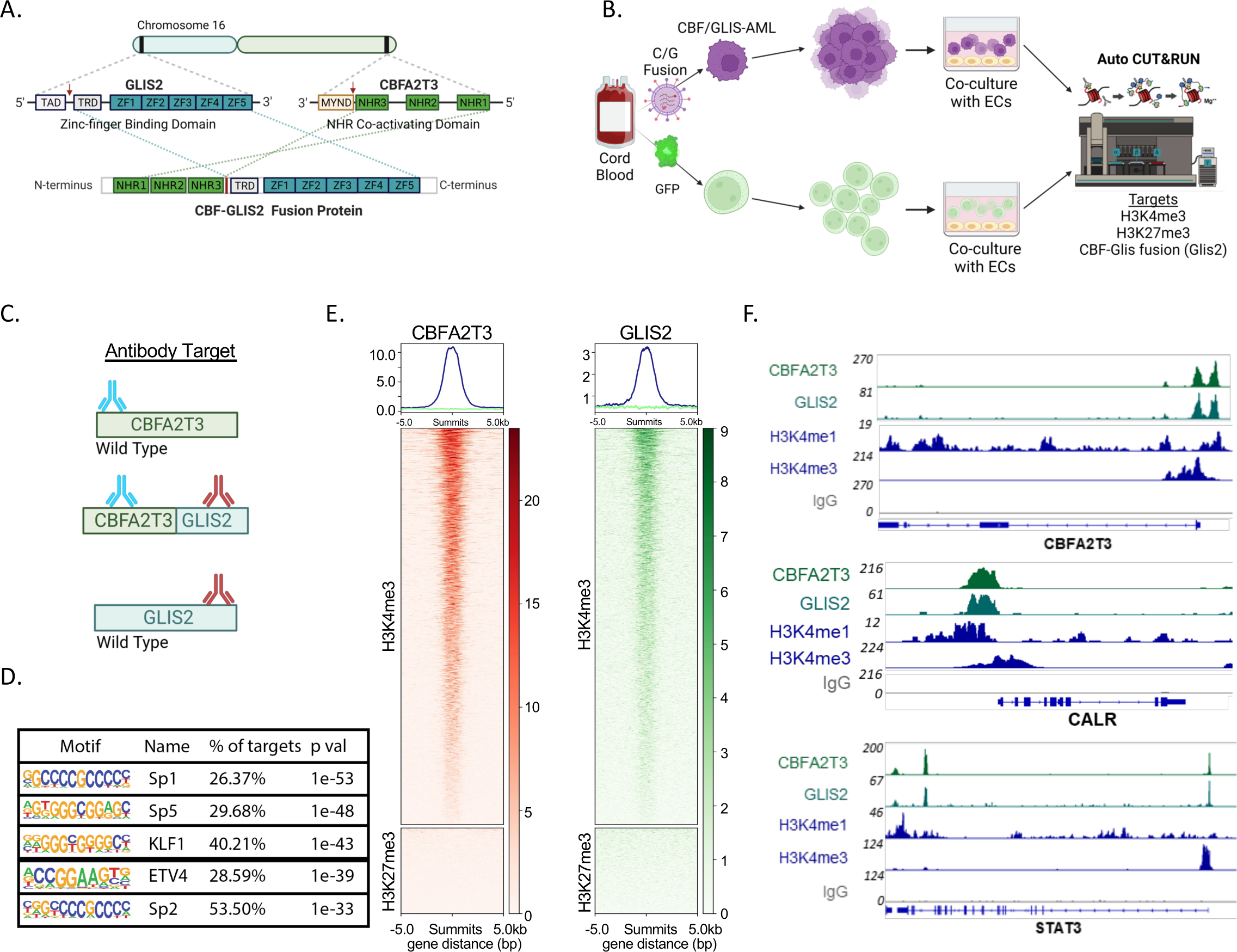
**A**: Schematic of the *CBFA2T3* and *GLIS2* locations on chromosome 16, protein domain map and resulting oncofusion protein with relevant domains. TAD: Transactivating domain; TRD: Transcriptional Repressor Domain; ZF: Zinc Finger Domain; MYND: *Myleoid, Nervy, Deaf-1* domain; NHR, Neutralized Homology Repeat Domain. Produced with *Biorender*. **B.** Schematic of in vitro leukemia system and experimental design using automated CUT&RUN assay. **C.** Schematic of antibodies targeting CBFA2T3 and GLIS2 used in CUT&RUN experiments. **D.** Motif analysis of consensus C/G peak set with motifs, transcription factor family name, % of peaks with motif and p-values. **E.** Average plots of CBFA2T3 and GLIS2 CUT&RUN signal plotted on H3K4me3 and H3K27me3 CUT&RUN peaks from C/G-transduced HSPCs. **F.** Integrated Genome Viewer (IGV) Browser Tracks of CUT&RUN signal at *CBFA2T3, CALR,* and *STAT3* loci show C/G fusion binding associated with H3K4me1 and H3K4me3 signal.

Here, we use Cleavage Under Targets and Release Using Nuclease (CUT&RUN)^5^ chromatin profiling technology to identify C/G fusion binding sites in an umbilical cord hematopoietic stem and progenitor (HSPC) model of C/G AML (Fig. 1b). We find C/G fusion targeting is determined by the Zf DNA binding domain of GLIS2, with CBFA2T3-mediated protein-protein interactions playing a less significant role in genomic localization. By comparing C/G binding sites to chromatin features in untransduced cord blood cells we identify a key subset of C/G targets that are normally silenced by Polycomb Repressive Complex 2 (PRC2) in HSPCs including *MYCN, ZFPM1 (FOG-1, Friend of GATA-1), ZBTB16 (PLZF, Promyelocytic Leukemia Zinc Finger Protein)* and *WT1 (Wilms Tumor 1).* Several of these genes are dependencies in genome-wide CRISPR screens in C/G AML cells. We show that untransformed megakaryocyte progenitors already express C/G fusion targets, suggesting that they may be particularly susceptible to transformation by the C/G oncoprotein. We identify the SMARCA4/BRG1 enzyme as a chromatin-related dependency in C/G AML and show that treatment with clinically-relevant BRG1 inhibitors potently kills C/G AML cell lines and extends survival in an aggressive xenograft model. BRG1 CUT&RUN profiling reveals that it binds regulatory elements near the *CBFA2T3* promoter and drives fusion expression, and BRG1 inhibitor treatment results in downregulation of the C/G fusion, revealing a novel therapeutic strategy in a pediatric AML subtype associated with poor prognosis.

## Results

### Identification of CBFA2T3-GLIS2 binding sites using CUT&RUN

To understand how the C/G fusion alters chromatin regulation to promote oncogenesis, we first sought to identify and characterize C/G binding sites in an umbilical cord blood model system of C/G AML^4^. We applied CUT&RUN technology^5^ to C/G-transduced cord blood cells using antibodies against both CBFA2T3 and GLIS2 (Fig. 1c). While the CBFA2T3 antibody has the potential to recognize both the wildtype CBFA2T3 protein as well as the fusion, the GLIS2 antibody recognizes the fusion only since wildtype *GLIS2* is minimally expressed in the hematopoietic system, as evidenced by transcriptomic analyses of GFP-transduced umbilical cord blood HSPCs, CD34+ PBSCs and normal bone marrow. (Fig. S1a)^2,3^. We previously validated this strategy for mapping oncogenic fusion protein binding sites in the context of KMT2A-rearranged leukemias, where the excellent signal to noise ratio of CUT&RUN allowed us to readily identify KMT2A fusion binding sites^6^.

To obtain a high-confidence set of C/G binding sites all samples were run in technical duplicates. Hierarchical clustering showed that replicates had the highest concordance, and that CBFA2T3 and GLIS2 samples from transduced cord blood cells were most similar to each other (Fig. S1b). The specificity of the GLIS2 antibody was further tested by performing CUT&RUN in untransduced cord blood cells, yielding minimal signal (Fig. S1c). To identify a stringent C/G peak set we merged replicates and performed peak calling using Sparse Enrichment Analysis for CUT&RUN (SEACR)^7^. This approach yielded 1843 overlapping peaks between CBFA2T3 and GLIS2 samples, and motif analysis on this peak set identified zinc finger-containing DNA binding proteins including SP1 and KLF1 as top hits. These results differ from a previous report that found the ERG/ETS motif as the dominant C/G motif target^8^ in the context of a C/G-GFP fusion overexpression study. However, the ETS-family member ETV4 motif was enriched in our dataset. Our motif analysis suggests that the Zf domains of GLIS2 are primarily responsible for the genomic localization of the C/G fusion in the umbilical cord HSPC model system.

We next mapped C/G binding sites onto H3K4me3 (active promoters) and H3K27me3 (facultative heterochromatin) sites in the C/G-transduced cells to determine whether the fusion localizes to active or repressed genes. Both CBFA2T3 and GLIS2 showed strong enrichment at active promoters (Fig. 1e), consistent with a role for the fusion in gene activation^8,9^. Minimal enrichment was observed at H3K27me3 sites (Fig. 1e). We also queried C/G binding sites identified in other ectopic expression studies and found strong signal at *CBFA2T3* and *CALR* loci (Fig 1f), indicating that the C/G fusion in our umbilical cord blood system binds loci shown to be C/G targets in other models^8^. Other loci previously shown to be relevant for C/G AML biology were also identified as C/G binding sites, including *STAT3* (Fig. 1f)^10^. Consistent with our genome-wide profiling, H3K4me1 and H3K4me3 peaks overlapped with C/G binding sites at these loci (Fig. 1f). We conclude that C/G fusion binding is associated with transcriptional activation of biologically relevant loci in the umbilical cord HSPC model system.

### Comparison of chromatin features in C/G-transduced cells vs controls

We explored how global chromatin landscapes in C/G- and GFP-transduced HSPCs relate by profiling H3K4me1, H3K4me3 and H3K27me3 in these two populations. Despite the strong leukemia phenotype in C/G-transduced cells and the inability of control HSPCs to proliferate robustly in vitro we found these chromatin features to be quite similar overall, with active enhancers and promoters showing similar enrichment profiles (Fig. 2a, Fig. S2a). In addition, 5487 H3K4me3 peaks were shared between samples, with an additional 921 being unique to GFP control cells and 1666 specific to C/G cells (Fig. S2b). Facultative heterochromatin sites as demarcated by H3K27me3 were also largely conserved between C/G and GFP-transduced HSPCs (Fig. 2a). This finding suggests that instead of massively reprogramming HSPC epigenomes the C/G fusion subtly modulates the regulatory element landscape to drive expression of a small number of key targets involved in oncogenesis.

**Figure 2:**
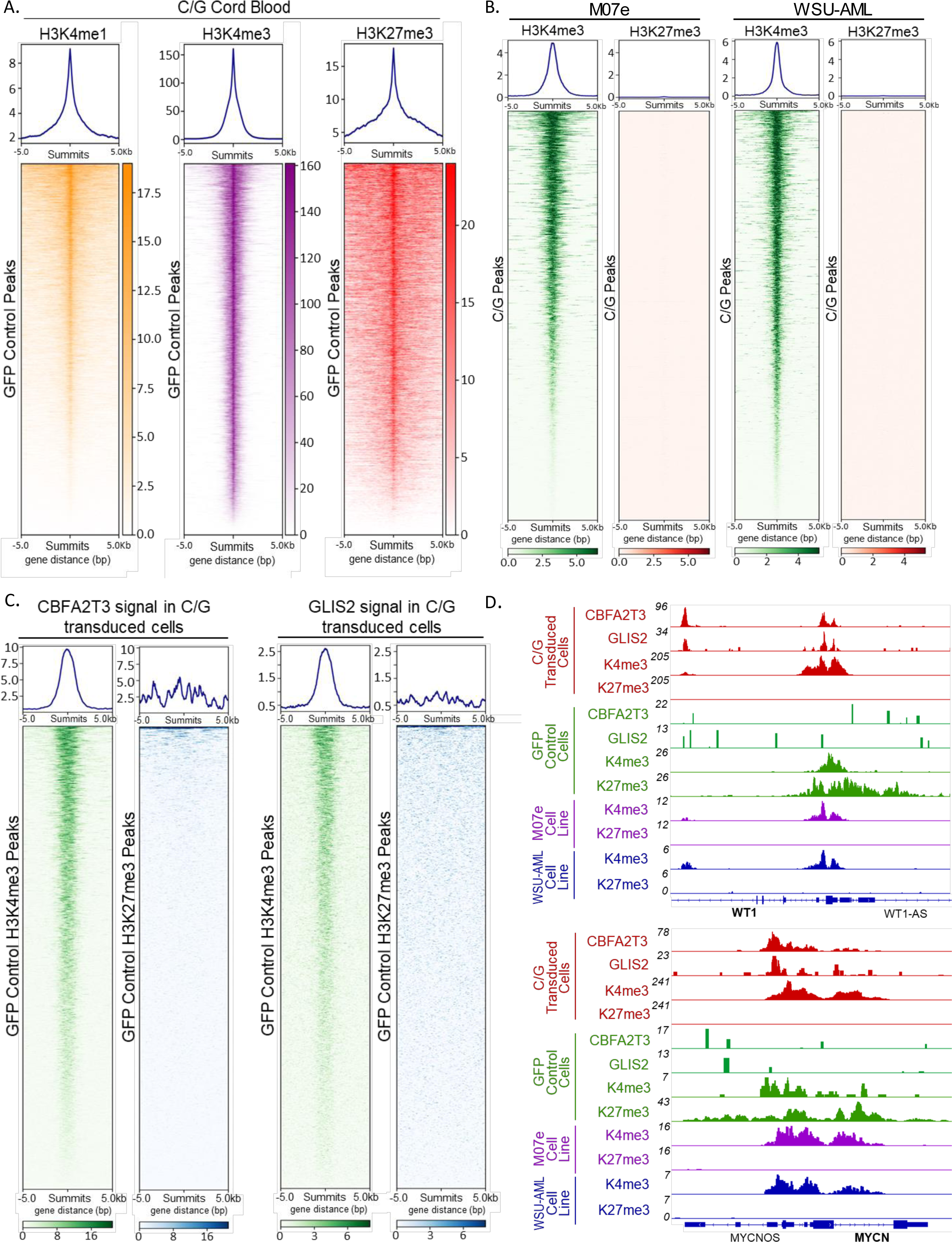
**A.** Average plots and heatmaps of H3K4me1, H3K4me3 and H3K27me3 CUT&RUN signal from C/G-transduced cells plotted onto H3K4me1, H3K4me3 and H3K27me3 peaks, respectively, from GFP-control cells. **B.** Average plots and heatmaps of H3K4me3 and H3K27me3 CUT&RUN signal from M07e and WSU-AML cell lines plotted onto consensus C/G CUT&RUN peaks from C/G-transduced cells. **C.** Average plots and heatmaps from CBFA2T3 and GLIS2 CUT&RUN signal in C/G-transduced cells plotted on H3K4me3 and H3K27me3 peaks in control HSPCs. **D.** Integrated Genome Viewer (IGV) Browser Tracks of CUT&RUN signal at *WT1* and *MYCN* loci show C/G fusion binding in C/G-transduced cells and H3K27me3 domains in GFP-control cells. H3K4me3 and H3K27me3 signal from M07e and WSU-AML cell lines are also shown.

Because of the similarity in active and repressed chromatin landscapes between control and C/G-transduced HSPCs we wondered whether C/G patient-derived cells shared these same chromatin features. We profiled H3K4me3 and H3K27me3 modifications in two patient-derived cell lines, M07e and WSU-AML^4,8,9,11^ for comparison with our umbilical cord blood HSPC datasets. We found that H3K4me3 signal in the two patient-derived cell lines was enriched at C/G-bound loci in HSPCs whereas H3K27me3 did not show similar enrichment (Fig. 2b). This finding indicates that fusion binding sites in C/G-transduced cells are associated with active chromatin features in patient-derived cell lines. Hierarchical clustering revealed that H3K4me3 profiles between the two transduced HSPCs and two patient-derived cell lines had similar correlation coefficients (Fig. S2a). In contrast, WSU-AML showed a more distinct K27me3 profile compared to the other samples (Fig. S2a). The similarity between H3K4me3 profiles across these samples is consistent with previous results showing that C/G cord blood-derived HSPCs recapitulate the transcriptome and immunophenotype of C/G patient specimens^4^.

To enrich for genes most impacted by C/G fusion binding in the cord blood HSPC system we compared C/G binding sites in transduced cells to H3K4me3 and H3K27me3 peaks in control HSPCs. Consistent with our comparison of H3K4me3 enrichment in C/G- and GFP-transduced cells, we found enrichment of CBFA2T3 and GLIS2 signal at H3K4me3 peaks in GFP-transduced HSPCs (Fig. 2c). Minimal overlap was observed between C/G binding sites in C/G-transduced cells and H3K27me3 domains in GFP control cells (Fig. 2c). Since PRC2 is critical for developmental hematopoiesis, we explored this list of 95 genes and found it contained factors important in AML, other cancers, and renal development including *MYCN*, *LMO2, ZBTB16 (Promonocytic Leukemia Zinc Finger, PLZF), Wilms Tumor 1 (WT1)* and *ZFPM1 (Friend of GATA, Family Member 1, FOG-1)* (Fig. 2d, Table 1). Examination of K4me3 and K27me3 signal at these loci in M07e and WSU-AML cells shows that they possess strong H3K4me3 signal at promoters and lack H3K27me3, consistent with gene activation. We conclude that the C/G fusion drives activation of a subset of key genes that are PRC2 targets in control HSPCs.

### C/G Targets in Umbilical Cord HSPCs are Expressed in C/G AML Patients

To investigate the relationship between C/G binding sites in the cord blood system and gene expression in C/G AMLs we integrated the 1843 C/G peaks with a comprehensive dataset from pediatric AML patients^10^. RNA-seq data from normal bone marrow specimens and GFP-control cells were also included in this analysis, as were 1-, 6- and 12-week timepoints post-C/G fusion transduction^4,11^. Of the 1616 peaks, that overlapped with an annotated gene, 1359 overlapped with genes that were not differentially upregulated in C/G specimens compared to normal bone marrow specimens or other AMLs (Fig. 3a). However, 111 overlapped genes that were differentially upregulated compared to normal bone marrow specimens, 51 overlapped genes that were upregulated in C/G AMLs compared to other AMLs, and 95 overlapped genes that were upregulated compared to both other AMLs and to normal marrow (Fig. 3a). Focusing on genes that are fusion targets but marked by H3K27me3 in control cells, we assessed *MYCN, ZBTB16, and ZFPM1* expression in patient samples. All three genes had higher median TPMs (Transcripts per Million) in C/G-transduced cord blood cells than control cells at the week 6 timepoint (Fig. 3b,c,d). They were also more highly expressed in C/G-positive AML specimens than in C/G-negative AMLs (Fig. 3b,c,d). In addition, comparision of expression levels of the 95 C/G targets marked by H3K27me3 in control cells found that thse genes are upregulated in a similar pattern in patient samples and cord blood cells at week 6 and 12 timepoints (Fig. S3).

**Figure 3.**
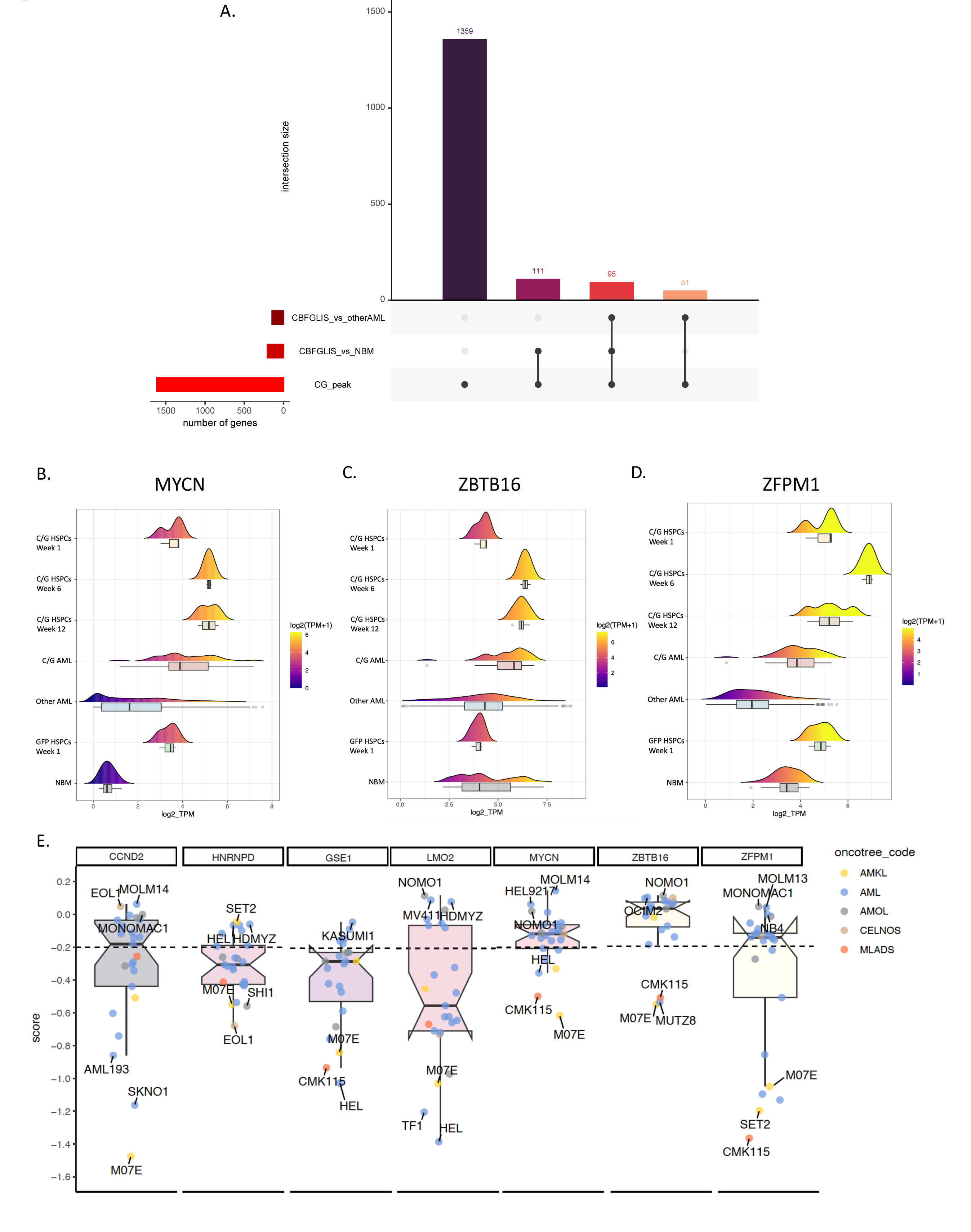
**A.** UpSet plot showing the overlap of upregulated differentially expressed genes from the comparison of “CBFGLIS patients vs other AML”, and “CBFGLIS patients vs normal marrows (NBM”) with peaks bound by C/G fusion. **B.** Density plot of *MYCN* expression from bulk RNA-seq dataset including C/G-transduced HSPCs at 1, 6, and 12 week timepoints, C/G AML patient samples, non-C/G AML patient sample, GFP-transduced HSPCs at 1 week time point and normal bone marrow. **C.** Same as B. except for *ZBTB16.* **D.** Same as B. except for *ZFPM1.* **E.** Box plot of DepMap dependency scores for key genes identified as C/G fusion targets in C/G-transduced HSPCs and possessing H3K27me3 domains in control HSPCs in myeloid-origin cell lines. AMOL, acute monocytic/monoblastic leukemia; CELNOS, Chronic Erythrocytic Leukemia Not Otherwise Specified; MLADS, Myeloid Leukemia Associated w/Down Syndrome.

These data indicate that key C/G targets in the HSPC system are upregulated in patient samples.

Within the C/G patient data we also noted that while *MYCN, ZBTB16, ZFPM1* are expressed in C/G positive and negative AMLs (Fig. 3b,c,d), almost all C/G patients displayed >2 TPM expression of these three genes. The finding that C/G AML patient samples show upregulation of these key genes while non-C/G AMLs can lack expression suggests that C/G AMLs may preferentially rely on these genes for proliferation. To explore the relationship between the 95 C/G targets that have H3K27me3 domains in control cells and growth we queried this list in the Cancer Dependency Map (DepMap) database^12,13^ of CRISPR dependency screens^12,13^. Out of the 24 myeloid leukemia cell lines in the database the lone C/G cell line, M07e, has the most negative *MYCN* dependency score at -0.7 and *ZBTB16* dependency score at -0.6 (Fig. 3e). *CCND2* displayed the most selective dependency score of -1.6 compared to median AML score of -0.2 (Fig. 3e). *LMO2* and *ZFPM1* also displayed strong negative dependency scores (Fig. 3e). This analysis demonstrates a functional consequence to expression of these factors, in contrast to many other AML subtypes where disruption of these factors does not impair proliferation (Fig. 3e).

### C/G Targets Are Expressed in Differentiating Umbilical Cord HSPCs

We also noted that at the week 1 timepoint many C/G targets are not upregulated in the cord blood system (Fig. S3), in contrast to 6 and 12 week timepoints (Fig. 3b,c,d) despite relatively efficient lentiviral transduction^4^. We hypothesized that a small population of cells within the cord blood pool is particularly susceptible to transformation by the fusion and takes several weeks to grow out, resulting in delayed detection of the C/G fusion signature in bulk RNA-seq data. To identify a population of sensitive cells we performed single cell RNA-seq on untransduced umbilical cord blood cells cultured in media supplemented with myeloid-promoting cytokines for 4 days, inducing HSPC differentiation (Fig. 4a). Using established markers and gene signatures (Fig. S4a), we identified HSCs, multipotent/common myeloid progenitors (MPP/CMP), granulocyte/monocyte precursors (GMP) and megakaryocyte/erythroid precursors (MEP) (Fig. 4b).

**Figure 4:**
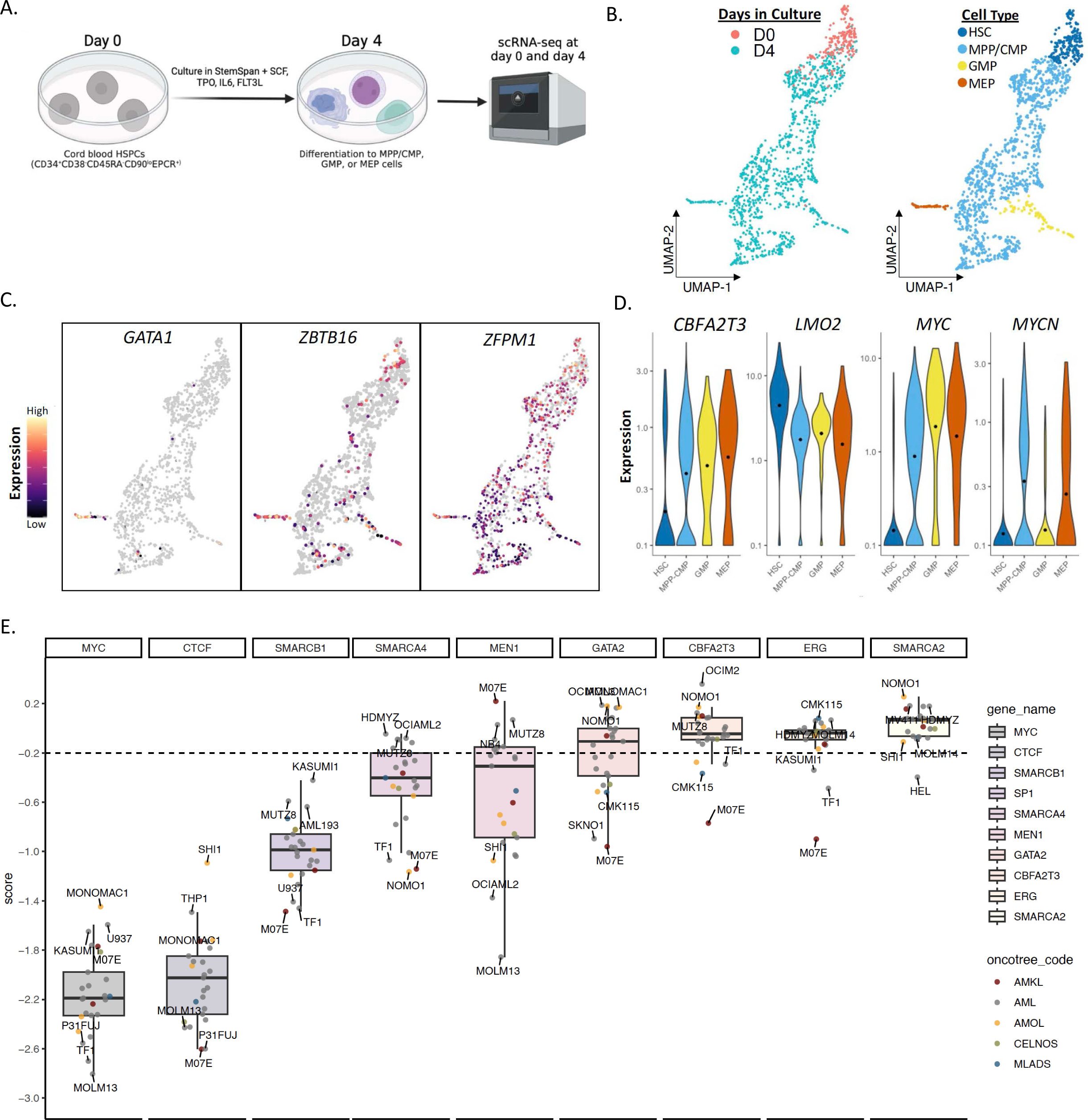
**A.** Schematic representation of single cell RNA-seq experiment in umbilical cord blood HSPCs and UMAP representation of annotated clusters. **B.** UMAP annotated by imputed cell type. MPP/CMP = multipotent progenitors/common myeloid progenitors, MEP = Megakaryocyte/erythroid progenitors, GMP = granulocyte/monocyte progenitors. **C.** UMAP representation of gene expression for indicated genes. **D.** Violin plots of gene expression (transcripts per million) for labeled genes by cell type. **E.** Box plot of DepMap dependency scores for key genes. AMOL, acute monocytic/monoblastic leukemia; CELNOS, Chronic Erythrocytic Leukemia Not Otherwise Specified; MLADS, Myeloid Leukemia Associated w/Down Syndrome.

We next explored several C/G targets and other key genes in this dataset. *GATA-1* was previously shown to be expressed in C/G AML^8^, is expressed in megakaryocyte precursors and showed highest expression in the MEP-annotated cluster (Fig. 4b,c). *MYCN* showed higher expression in MEPs than GMPs or HSCs (Fig. 4d), in contrast to *MYC,* which was expressed at similar levels in MPP/CMPs, GMPs, and MEPs (Fig. 4D, Fig S4b). *ZBTB16 and ZFPM1* expression were higher in MEPs than GMPs and MPPs (Fig. 4c). In contrast, *LMO2* was expressed in all clusters. We also explored *GATA-2* expression as it interacts with ZFPM1 and plays a critical role in megakaryocyte development, and it was expressed at higher levels in MEPs than other populations. Intriguingly, *GATA-2* is also a top 10 selective dependency in the M07e cell line in DepMap (Fig. 4e). Though *CBFA2T3* is also a selective dependency (Fig. 4e), its expression was not restricted to MEPs (Fig. 4d). Preferential expression of *ZBTB16, ZFPM1, GATA-2* and *MYCN* in MEPs may help to explain the association between the C/G fusion and an AMKL immunophenotype^11^. These data show that while MEPs express an optimal confluence of factors to permit context-specific leukemogenesis by the C/G fusion, other HSPC populations can also express key genes, providing insight into how non-megakaryocytic C/G AMLs can arise.

### SMARCA4/BRG1 Inhibition as a Therapeutic Vulnerability in C/G AML

Due to the poor prognosis associated with C/G AML, we sought to capitalize on our chromatin profiling and transcriptomic analyses to identify therapeutic strategies. Menin *(MEN1)* is a critical chromatin-related leukemogenic factor in several AML subtypes^14^ and clinically-tractable inhibitors have shown efficacy in recent trials^15^. We wondered whether menin may play a role in C/G AML. Examination of DepMap found that the M07e cell line has the most positive dependency score of all myeloid-derived cell lines in the database despite being expressed at levels similar to some KMT2Ar cell lines (Fig. 4e, S, where it is a strong dependency. This finding indicates that C/G AMLs do not depend on menin-related functions for proliferation.

We next explored the mammalian Brahma Associated Factor (BAF) complex, a member of the Switch/Sucrose Non-Fermenting (SWI/SNF) chromatin remodeler family, given its role in antagonizing Polycomb Repressive Complex activity during development^16^. We found that out of all AML cell lines in the DepMap database M07e has the second strongest dependency score for *SMARCA4/BRG1,* the ATPase subunit of the canonical BAF complex, at -1.06 (Fig. 4e, S4c), and third most negative score for cofactor *SMARCB1/BAF47* at -1.39 (Fig. 4e, S4c).

Encouraged by these dependency scores, we tested two highly selective, orally bioavailable inhibitors of the related ATPases BRG1/BRM, BRM-014 and FHD-286 in M07e and WSU-AML cell lines^17,18^. Both cell lines demonstrated sensitivity to these agents, with FHD-286 displaying IC50’s of 5nM in WSU-AML and 49nM in M07e and BRM-014 showing IC50s of 22nM in WSU-AML and 117nM in M07e (Fig. 5a). SMARCA2/*BRM* has a dependency score of 0 in M07e (Fig. 3e), supporting BRG1 and not BRM inhibition as the relevant FHD-286 target in C/G AML cell lines.

**Figure 5:**
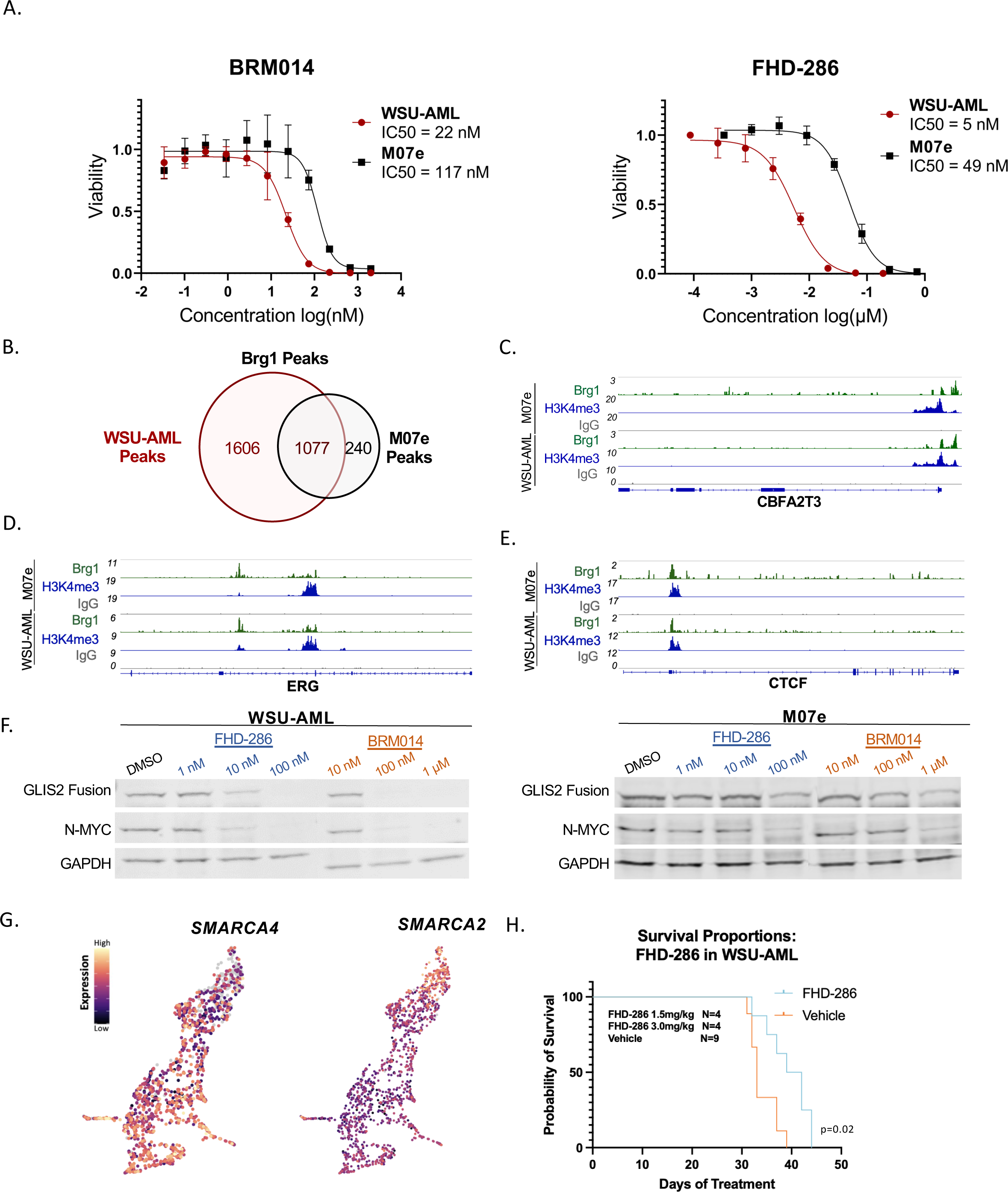
**A.** Viability curves (Cell Titer Glo Assay) for BRG1/BRM inhibitors in M07e and WSU-AML cell lines treated for 72hrs. **B.** Number of overlapping and unique BRG1 CUT&RUN peaks in M07e and WSU-AML cell lines. **C-E.** IGV browser visualization of key BRG1 peaks with associated H3K4me3 signal. **F.** Western blotting of C/G fusion (GLIS2 antibody), N-MYC and GAPDH in WSU-AML and M07e cells treated with BRG1/BRM inhibitors for 48hrs. **G.** UMAP representation of *SMARCA4/BRG1* and *SMARCA2/BRM* expression in scRNA-seq dataset from untransduced HSPCs. **H.** Survival curves for WSU-AML xenograft model comparing vehicle and FHD-286-treated mice.

To understand the chromatin-level effects of BRG1 inhibition in C/G AML cells we mapped BRG1 binding sites in untreated M07e and WSU-AML cells using CUT&RUN. We identified 1606 peaks in the WSU-AML cell line and 1247 in M07e (Fig. 5b). These peak sets had substantial overlap, with 1077 genes in common between the two cell lines. We then integrated this peak set with M07e DepMap data to find BRG1-bound genes that had strong dependency scores. This analysis yielded interesting genes including *CBFA2T3*, *ERG* and *CTCF* (Fig 4e, 5.c,d,e) as well as several splicing factors including *SRSF2* and *SRSF3* (Table 2). *CBFA2T3* is the 5’ partner of the C/G fusion event and the promoter is used to drive expression. The mesenchymal transcription factor *ERG* was previously shown to be critical for C/G leukemogenesis^8^. *CTCF* and alternative splicing factors including *SRSF2* and *SRSF3* are involved in leukemogenesis in other myeloid-derived malignancies^19,20^, but do not have established roles in C/G AML.

The enrichment of BRG1 at the *CBFA2T3* locus suggests that BRG1 inhibition will reduce C/G fusion expression. We performed Western blotting for the C/G fusion in WSU-AML and M07e cells treated with FHD-286 and BRM-014 and found substantial reductions in levels (Fig. 5f). We also explored N-MYC levels since the *MYCN* locus is a direct C/G fusion target but does not have a BRG1 peak in CUT&RUN data and found that levels were reduced. In addition, several other BRG1 targets not bound by the C/G fusion including CTCF and ERG were also reduced by the BRG1 inhibitors (Fig. S5b). The dependency of these factors on BRG1 suggests that C/G AMLs may arise from a cell of origin that is particularly dependent on *SMARCA4* for developmentally-regulated gene expression. To test this, we queried *SMARCA4* expression in our scRNA-seq dataset and found higher expression in MEPs and GMPs than HSCs and MPPs (Fig. 5g), whereas *SMARCA2* is preferentially expressed in HSCs (Fig. 5g). These findings are consistent with previous studies showing *SMARCA4* and SMARCA2 play non-overlapping roles in hematopoiesis^21^. In AML samples *SMARCA4* is not significantly upregulated in C/G patients compared to other AMLs (Fig. 5c), though a subset of C/G patients do display increased *SMARCA4* expression compared to other AMLs, CD34+ peripheral blood HSPCs and normal bone marrow.

Given the encouraging results in vitro we tested FHD-286 in an aggressive murine xenograft model derived from luciferase-labeled WSU-AML cells. Since 100% of mice injected with WSU-AML develop leukemias and succumb to disease within 45 days of injection, we initiated FHD-286 dosing 7 days after cell line injection at 1.5mg/kg daily orally. Few studies have reported on use of this agent in preclinical models so we monitored mice for 1 week before increasing a subset of the cohort to 3 mg/kg/dy. The median survival for vehicle mice was 33 days and for all FHD-286 treated mice was 40.5 days (Fig. 5h). Subgroup-specific analyses found that the 1.5mg/kg group had a median survival of 43 days while the 3mg/kg/dy group’s median survival was 38 days (Fig. S5d). Leukemic burden as assessed by luminescence did not appear to be affected by FHD-286 despite the prolonged survival (Fig. 5e). We conclude that BRG1 inhibition is a viable therapeutic strategy for C/G AML.

## Discussion

Transcription factor fusion proteins occur in a diverse array of cancers and are often the sole event underlying transformation. Despite their immense oncogenic potential, they display tremendous context specificity and are usually restricted to specific cancer subtypes. Here we capitalize on an in vitro transformation system to understand how the C/G fusion cooperates with a permissive chromatin landscape to drive a lethal infant AML subtype. By profiling C/G binding sites in an umbilical cord blood-derived HSPC model system, we find that the C/G fusion binds a small number of key loci instead of massively rewiring the epigenome. Integration of PRC2 sites in untransformed umbilical cord HSPCs with C/G fusion binding sites in leukemia-like cells identified *MYCN* as a target, indicating that MYC-family members play a more significant role in C/G AML than previously appreciated. *ZBTB16, ZFPM1* and *LMO2* are also C/G targets and DepMap dependencies, and may comprise a core regulatory circuit along with *GATA-1* and *GATA-2* in this subtype of AML as *LMO2* does in other AMLs^18^. Single cell transcriptomic analyses of differentiating, untransduced cord blood cells found that many of these genes are already expressed in megakaryocyte progenitors, and that *ZBTB16* and *ZFPM1* are preferentially expressed in MEPs and GMPs. These findings help to explain the association of the C/G fusion protein with an AMKL phenotype^11^.

By integrating epigenomic and transcriptomic data with genome-wide CRISPR screens we identified the chromatin remodeler *BRG1*/*SMARCA4* as a critical dependency in C/G AML. Profiling of BRG1 binding sites in C/G leukemia cells shows that it binds and drives expression of the C/G fusion along with other C/G AML dependencies including *ERG, SRSF2,* and *CTCF*. Treatment with clinically-tractable BRG1 inhibitors found that they reduce C/G fusion levels as well as levels of downstream targets, induce apoptosis as measured by cPARP (Fig. S5b), and extend survival in murine xenografts. Single cell analyses show that *SMARCA4* is upregulated in MEPs, suggesting that gene expression in this subset of HSPCs may be particularly dependent on *SMARCA4/BRG1* and other BAF components. This is in contrast to HSCs that may rely more on *BRM/SMARCA2* remodeling activity.

Although much insight into C/G leukemogenesis was gained by these experiments, new directions were also identified. For example, despite M07e and WSU-AML cells both possessing the C/G fusion, WSU-AML cells are 4-fold more sensitive to BRG1 inhibition than M07e. The reason for this differential sensitivity is unclear, though M07e has a secondary NRAS mutation that is not normally seen in C/G AML samples that may have been acquired to facilitate growth *in vitro*^22^. Additional patient-derived cell lines and murine models are needed to investigate this difference and determine whether C/G AML patients may respond differently to BRG1 inhibition. In addition, AML cell lines display substantial variation in sensitivity to BRG1-targeting gRNAs in DepMap, being a critical dependency in some and neutral in others. This has also been observed in the context of BRG1 inhibitors^17,18^. The sensitivity to BRG1 inhibition has been ascribed to *PU.1 (Sp1)*^17,18^, however, in DepMap *PU.1* is a neutral gene in M07e. Further, we observed apoptosis in response to BRG1 inhibition, while differentiation has been reported in other AMLs^18^. Thus, other BRG1 targets are likely mediating the sensitivity in C/G AML, including the C/G fusion itself. It is also possible that BRG1 inhibition may have additional targets in other AML subtypes beyond PU.1.

The dependency of C/G AML on *MYCN* is intriguing. Many leukemias including C/G AML are dependent on C-MYC (Fig. 4e), though M07e is less dependent on C-MYC than most other AML cell lines (Fig. 4e, S4c). Whether N-MYC expression accounts for this finding and a role for it in hematological malignancies is less clear. Further, few cancers express and rely upon both factors for proliferation. Previous work has shown that *MYC* and *MYCN* knockout mice have different hematopoiesis-specific defects, and the dual knockout has a distinct phenotype from either single knockout^23^. How these two factors cooperate to drive C/G leukemias remains unknown, but a previous report showing that C/G AMLs are sensitive to aurora kinase inhibitors^24^ may in part be explained by N-MYC being an important an aurora kinase substrate^25^.

The dependency on CTCF is also noteworthy, as M07e has the most negative dependency score for this factor among all leukemias in the DepMap database. A recent report identified a *CBFA2T3-CTCF* fusion in a patient with relapsed AML and showed that this fusion upregulated STAT3 expression, a direct C/G fusion target in our cord blood system^26^. Whether C/G AMLs are particularly dependent upon CTCF-mediated local effects such as insulation or long-range contacts merits further investigation.

Recent studies have shown that BRG1 inhibition may be a tractable therapeutic strategy in AML, and a clinical trial with FHD-286 is currently underway in adult AML. Our work suggests that certain AMLs may be preferentially dependent on BAF-dependent chromatin remodeling, either to drive expression of key oncogenic factors such as fusion proteins or maintain expression of dependencies such as CTCF and splicing factors. Profiling of BRG1 and CTCF sites in C/G leukemias and other AMLs, along with 3D-interaction mapping methods such as Hi-C may reveal AML subtype-specific roles for these factors. Finally, as new immunotherapies and targeted approaches for C/G AML are developed, the impact of epigenetic therapies on target expression such as *FOLR1* in C/G AML^4^ must be investigated as combination strategies are needed to treat these refractory malignancies.

## Acknowledgements

The authors acknowledge Research Scientific Computing at Seattle Children’s Research Institute for providing HPC resources. We thank Foghorn Therapeutics for providing FHD-286 for murine trials. The Fred Hutchinson Cancer Center Genomics Core including Phil Corrin and Feinan Wu are acknowledged for their assistance with sample preparation and data analysis. Jorja Henikoff, Terri Bryson, and Christine Codomo are acknowledged for their assistance with sample preparation, sequencing and data analysis and Derek Janssens for helpful discussions. We thank Lisa Ang and Heather Conti for assistance with murine xenograft studies. J.F.S is supported by a Hyundai Hope on Wheels Young Investigator Award and NCI 1K08CA256167.

## Methods

### RNA-sequencing Library Construction

Methods were reported previously in Quy et al. (https://www.jci.org/articles/view/157101#SEC4). In brief, total RNA was extracted for primary diagnostic pediatric AML samples, primary healthy normal bone marrow (NBM) controls, and CBFA2T3-GLIS2 transduced cord blood (C/G-CB) and GFP control (GFP-CB) cells in endothelial coculture^4^. Paired-end mRNA libraries (75-bp strand-specific) were constructed using the ribodepletion v2.0 protocol by the British Columbia Genome Sciences Center (Vancouver, BC, Canada). Reads were pseudo-aligned and quantified with Kallisto v0.45.0 using a GRCh38 reference. Gene-level counts were normalized to transcripts per million (TPM) using tximport v1.16.1.

### Transcriptome Analysis

Differentially expressed genes were identified using limma voom v3.50.3 with TMM normalized gene counts (EdgeR v3.36.0). Differentially expressed genes with absolute log2 fold-change > 1 and Benjamini-Hochberg adjusted p-values < 0.05 were included for further analysis.

Unsupervised hierarchical clustering was carried out with ComplexHeatmap v2.10.0 using Euclidean distances with the ward.D2 linkage algorithm.

### Data availability

RNA-sequencing data for primary AML patients and healthy bone marrows can be accessed from the Database of Genotypes and Phenotypes (https://www.ncbi.nlm.nih.gov/gap/) under accession number phs000465.v19.p8 with a data use agreement. RNA-sequencing data from engineered CB are deposited in the NCBI Gene Expression Omnibus under accession GSE181726. RNA-sequencing counts matrices can be retrieved from the TARGET AML project under mRNA-seq DCC open access (https://target-data.nci.nih.gov/). Cut&Run data from human umbilical cord blood samples and cell lines are available on NCBI GEO at accession number GSE239849. Single-cell RNA-sequencing are in the process of being submitted to GEO.

### CUT&RUN Sequencing

Cut&Run data were generated from human umbilical cord blood samples transduced with CBFA2T3-GLIS2 or GFP control. Independent experiments were completed for biological duplicates. Details of transduction and cell culture conditions are reported in^4^. Cut&Run libraries were prepared at the Fred Hutchinson Genomics Core using the AutoCut&Run protocol and processed as reported in ^27^ using the GRCh38 reference genome. Sequencing reads were mapped to the human hg38 genome build using Bowtie2^28^. Regions of enrichment (peaks) were identified using SEACR v1.4 in stringent mode and IgG control normalization^7^. Peaks identified in both biological replicates were considered high confidence peaks and were retained for further analysis. Heatmaps were generated using deepTools version 3.5.1 ^29^. CBFA2T3-GLIS2 (C/G) peaks were identified by intersection of the high confidence peaks from CBFA2T3 and GLIS2 from C/G transduced cord blood samples using Bedtools intersect v2.30.0. Motif enrichment and peak annotations were completed using Homer version 4.11 with hg38 promoter sets and Homer motif files^30^.

Associations of CUT&RUN with transcriptome sequencing and DepMap data were carried out in the R 4.1 statistical environment and scripts are available by request. Gene effect scores from CRISPR knockout screens were downloaded from The Cancer Dependency Map Portal (https://doi.org/10.6084/m9.figshare.21637199.v2 and https://doi.org/10.6084/m9.figshare.22765112.v2). Peak annotations were de-duplicated and associated to transcriptome gene counts and DepMap scores using ensembl gene_ids, NCBI refseq IDs, and gene symbol. C/G peak annotations found to have H3K27me3 histone marks in GFP controls were identified using the union of regions in both H3K27me3 GFP replicates; these gene annotations were further investigated for DepMap gene dependencies in myeloid lineage cell lines and those with scores < -0.1 in M07e (C/AML cell line) were retained.

### Sorting

Human umbilical cord blood (CB) samples were obtained from normal deliveries at Swedish Medical Center (Seattle, Washington, USA). Fresh CB samples were processed with red blood cell lysis buffer, enriched for CD34^+^ cells using CliniMACS CD34 MicroBeads (Miltenyi Biotec, 130-017-501), and frozen down for the future use. On the day of sort, CD34^+^ cells from multiple pooled CB samples (8 units each) were thawed and stained with the following antibodies: CD45RA-APC-Cyanine7, CD38-PECy7 (Invitrogen), CD34-PerCP, CD90-BV421, EPCR-BV786, Lineage cocktail (CD3, 14, 16,19,20,56)-FITC (BD Biosciences) and DAPI (to exclude dead cells) and. Cells were sorted on a BD FACS Aria II equipped with BD FACS DIVA software as immunophenotypic HSC (DAPI^-^CD34^+^Lin^-^CD38^-^CD90^lo^CD45RA^-^EPCR^+^).

### Single cell RNA sequencing

For Day 0 (D0) CB HSC samples, ∼2,000 freshly sorted CD34^+^Lin^-^CD38^-^CD90^lo^CD45RA^-^EPCR^+^ cells were washed and resuspended with PBS containing 0.04% ultrapure BSA (Invitrogen) on ice for scRNAseq. For Day 4 (D4) samples, ∼2,000 CD34^+^Lin^-^CD38^-^CD90^lo^CD45RA^-^EPCR^+^ cells were cultured on tissue culture plates containing StemSpan II media (Stem Cell Technologies) supplemented with 50ng/ml recombinant human SCF (Miltenyi Biotec), recombinant human Flt3-ligand (Miltenyi Biotec), recombinant human IL-6 (Miltenyi Biotec), recombinant human TPO (PeproTech), and 10ng/ml recombinant human IL-3 (Gibco). Cultured cells were infected with lentivirus encoding GFP/Scrambled control shRNA overnight and fresh media was added the following day. Following a total of 4 days of culture, cells were harvested by pipetting, washed and resuspended with PBS containing 0.04% ultrapure BSA on ice for scRNAseq. Cell suspensions were loaded into the Chromium Single Cell B Chip (10X Genomics) and processed in the Chromium single cell controller (10X Genomics). The 10X Genomics Version 3 single cell 3’ kit was used to prepare single cell mRNA libraries with the Chromium i7 Multiplex Kit, according to manufacturer protocols. Sequencing was performed for pooled libraries from each sample on an Illumina NextSeq 500 using the 75 cycle, high output kit, targeting a minimum of 50,000 reads per cell.

### Single Cell Transcriptome Analysis

The Cell Ranger 2.1.1 pipeline (10X Genomics) was used to align reads to the GRCh38 reference genome and generate feature barcode matrix, filtering low-quality cells using default parameters. The Monocle3 (v.3.2.3.0) platform (https://doi.org/10.1038/s41586-019-0969-x)(https://doi.org/10.1038/nbt.2859) was used for downstream analysis, combining D0 and D4 samples for downstream analysis, using a negative binomial model of distribution with fixed variance, normalizing expression matrices by size factors. Batchelor (v.1.2.4) (https://doi.org/10.1038/nbt.4091) was used to remove batch effects between samples using a “mutual nearest neighbor” algorithm. Uniform Manifold Approximation (UMAP) was used for dimensionality reduction (https://doi.org/10.1128/mSystems.00691-21). Clustering was performed by Louvain method (https://doi.org/10.1016/j.cell.2015.05.047). Cell type classification was performed using the Garnett package (v.0.2.15) (https://doi.org/10.1038/s41592-019-0535-3) within Monocle3. Marker gene sets based on established cell type-specific genes (listed below) were used to train a classifier data and classify cell types in each cluster.

| Cell type | Genes expressed | Genes not expressed |
| --- | --- | --- |
| HSC | BEX1, GATA3 | CD33 |
| MPP-CMP | CD34, CD33 | GATA1, MPO, KLF1 |
| GMP | CEBPA, MPO, CD38 | CD34 |
| MEP | GATA1, KLF1 |  |

Gene set scores were calculated for each single cell as the log-transformed sum of the size factor-normalized expression for each gene in published signature gene sets^31,32^. R scripts used for analysis are available upon request.

### Cell Lines and Drug Treatments

The M07e cell line was seeded at 1x10^6^ cells/ml and cultured in RPMI (Gibco) +10%FBS +1X Glutamax (Gibco) + 10ng/ml IL-3 (STEMCELL Technologies). The WSU-AML cell line was also seeded at 1x10^6^ cells/ml, cultured in RPMI (Gibco) +10%FBS +1X Glutamax (Gibco). Both cell lines were treated with FHD-186 or BRM-014 for 48 hours. FHD-286 was added at 1nM, 10 nM, and 100 nM. BRM-014 was added at 10 nM, 100 nM, and 1000 nM. A DMSO control group was included with each drug treatment. For the CUT&RUN experiments, the M07e line was treated with 100 nM FHD-286 for 24 hours, while the WSU-AML line was treated with 10 nM FHD-286 for 24 hours.

### Western Blotting

Cells were lysed with 3% SDS lysis buffer. Lysates were separated using Tris-glycine gel electrophoresis then transferred to a 0.45 µM nitrocellulose membrane (Biorad). The membrane was incubated with primary antibodies, all diluted 1:1000, overnight at 4°C. The following primary antibodies were used: GLIS2 (Invitrogen, PA574829), N-MYC (Cell Signaling Technology, 84406S), and GAPDH (Cell Signaling Technology, 97166S). The secondary antibodies (LI-COR 800CW Goat anti-mouse and 680RD Goat anti-rabbit) were diluted 1:10,000 and incubated with the membrane for 1 hour at room temperature.

### Cell Viability Assay

The cell viability was performed in sterile 96-well clear-bottom plates with 200 ul of growth medium per well in triplicate. Viability was quantified with the CellTiter-Glo (Promega) assay and analyzed on a VICTOR X plate reader (Perkin Elmer).

### Animal Studies

NOD/SCID/γc−/− (NSG) mice were obtained from The Jackson Laboratory and housed at Seattle Childrens’ on a standard 12 hour light dark cycle with *ad libitum* food and water. All experiments were approved by Seattle Childrens’ Institutional Animal Care and Use Committee (ACUC00708) and in compliance with approved AAALAC standards.

Age-matched animals were randomly assigned to experimental groups by sex (with equal numbers of males and females in each arm). After tumor implant and throughout dosing, animals were monitored and scored for clinical symptoms of leukemia (tachypnea, anemia, and hind-limb paralysis) in addition to standard mouse health monitoring metrics (weight-loss, hunched posturing, piloerection). Moribund animals were humanely euthanized when any combination of tumor specific and/or standard health symptoms reached predetermined study criteria. For animal studies, 2 x 10^6^ eGFP-luciferase labeled WSU-AML cells were injected intravenously by tail vein injection. One week after tumor engraftment, animals were treated with either 1.5 mg/kg FHD-286 in 20% (2-Hydroxypropyl)-β-cyclodextrin in USP grade phosphate buffered saline or 20% H-β-cyclodextrin by oral gavage with a schedule of 5 days on and 2 days off. On day 8, FHD treated arm was randomly split into higher treatment 3.0 mg/kg or maintained at 1.5 mg/kg after finding the higher dose was well tolerated. Tumor burden was measured by bioluminescence imaging.

### Study approvals

De-identified human CB specimens used in this study were obtained after informed consent from maternal donors, following approval by the Swedish Medical Center Institutional Review Boards (de-identified research CB collected under Swedish IRB Protocol #5647; FHCC IRB non-human subjects research approval #6007-2167). The study was conducted in accordance with the Declaration of Helsinki.

## Supplementary Figures

**Figure S1:**
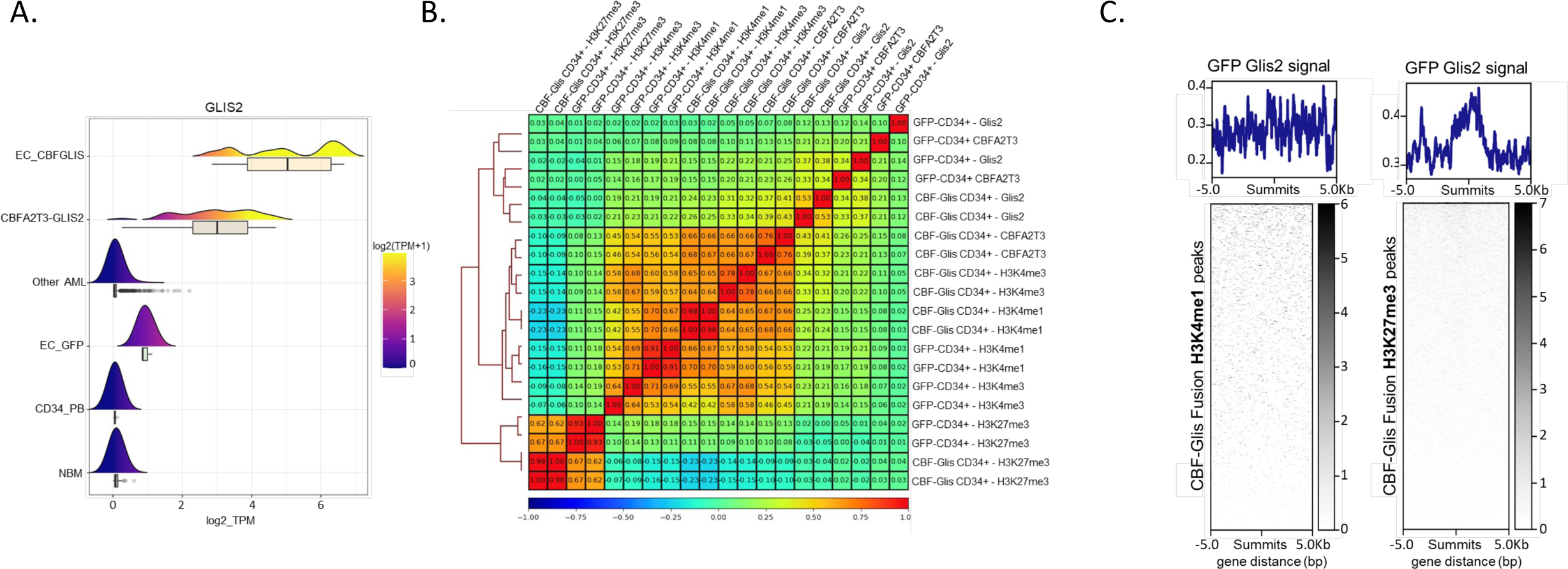
**A.** Density plot of *GLIS2* expression in a bulk RNA-seq dataset including C/G cord HSPC-endothelial cell co-culture system (EC_CBFGLIS), CBF-Glis patient samples (CBFA2T3- GLIS2), non-CG pediatric AML patients (Other AML), GFP control HSPCs cultured with endothelial cells (EC_GFP), CD34+ selected peripheral blood cells (CD34+_PB) and normal bone marrow specimens (NBM). **B.** Spearman correlation heatmap of CUT&RUN signal from umbilical cord HSPC samples. **C.** Average plots and heatmaps of GLIS2 CUT&RUN signal in GFP control HSPCs.

**Figure S2:**
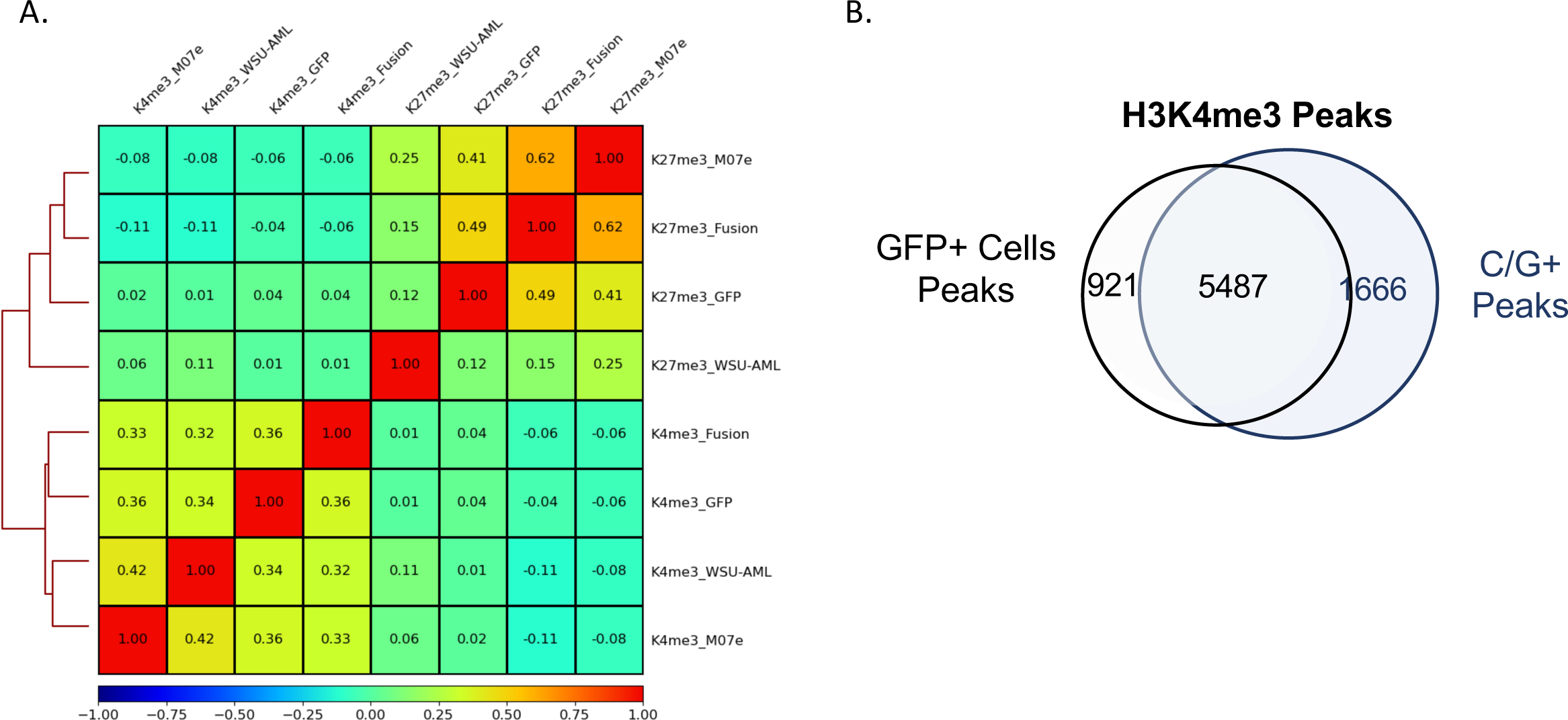
Spearman correlation heatmap of CUT&RUN signal with indicated antibodies from two C/G-patient derived cell lines and cord blood HSPCs transduced with either the C/G fusion or GFP control. **B.** Venn diagram showing overlap of H3K4me3 peaks in the umbilical cord blood model system.

**Figure S3:**
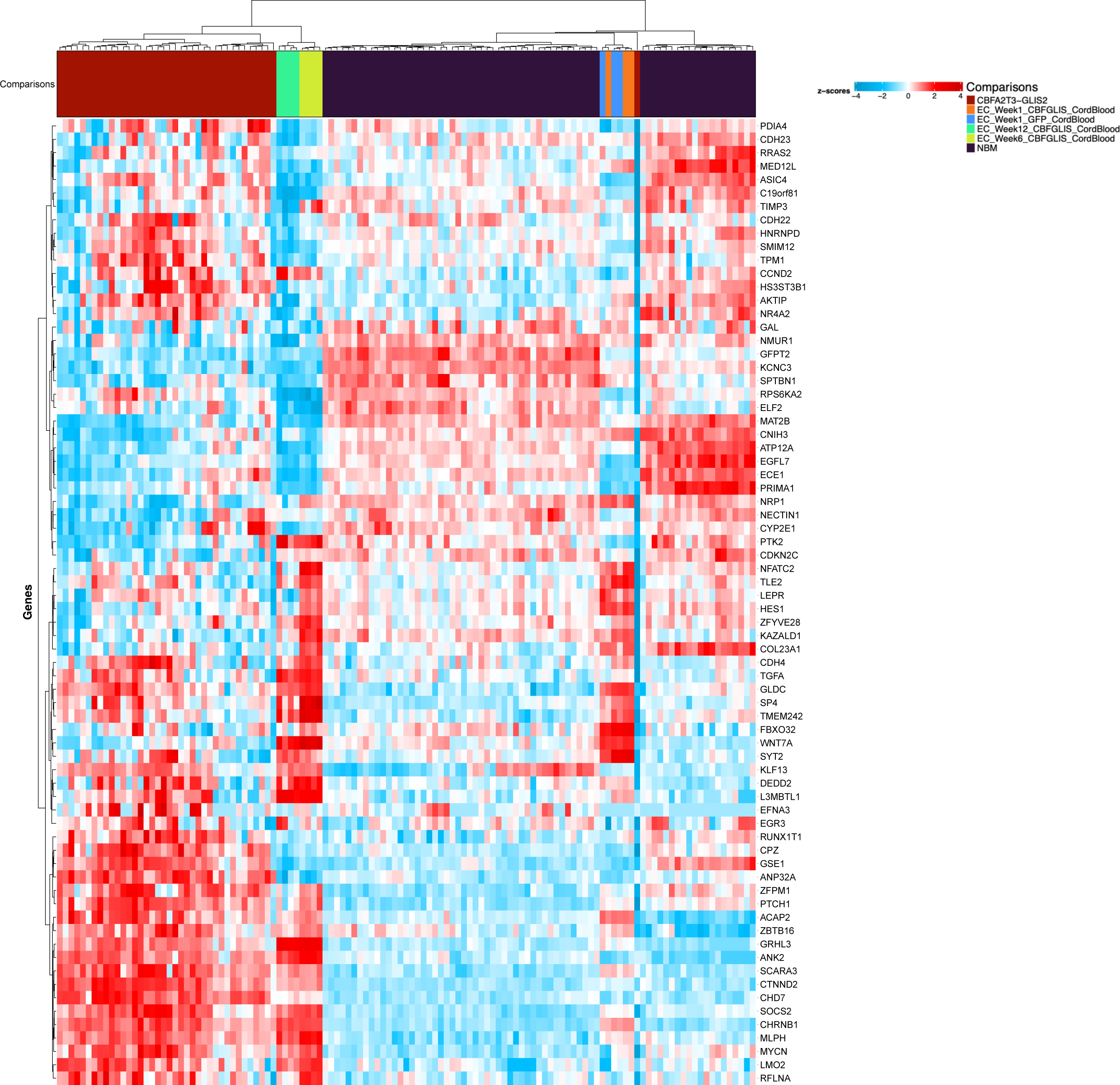
Heatmap of genes with C/G peak annotations and found to have H3K27me3 histone marks in GFP controls. Expression of these C/G gene targets is shown in a heatmap including primary C/G AML patient samples, C/G cord HSPC model and healthy controls. Expression values depicted are z-score transformed TPM normalized counts.

**Figure S4:**
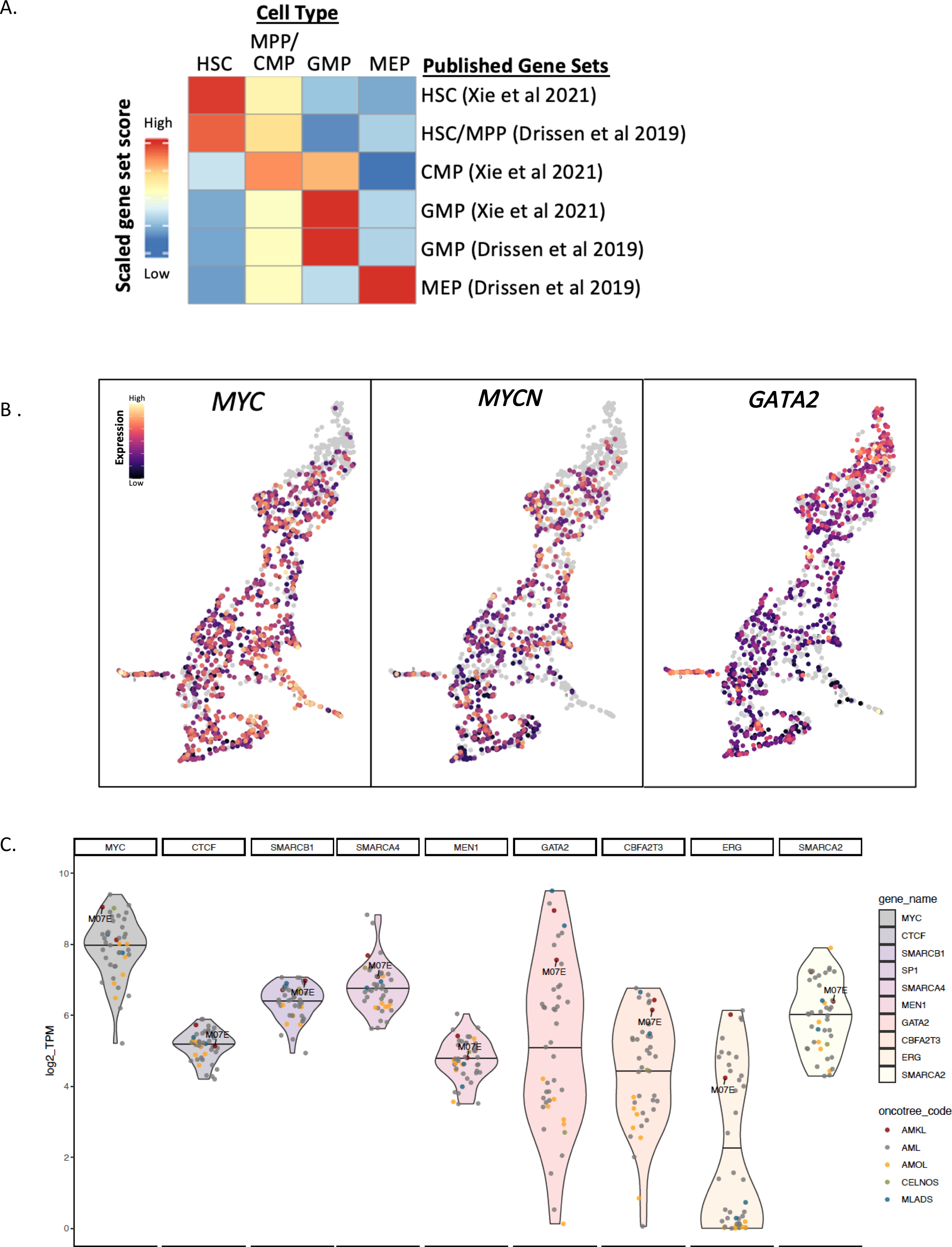
**A.** Heatmap of gene set scores by cell type for published gene sets associated with HSC, HSC/MPP, CMP, GMP, and MEP. **B.** Single cell gene expression of *MYC*, *MYCN* and *GATA-2* represented in UMAPs. **C.** Violin plots of gene expression (TPM) for key genes from the Cancer Cell Line Encyclopedia. AMOL, acute monocytic/monoblastic leukemia; CELNOS, Chronic Erythrocytic Leukemia Not Otherwise Specified; MLADS, Myeloid Leukemia Associated w/Down Syndrome.

**Figure S5:**
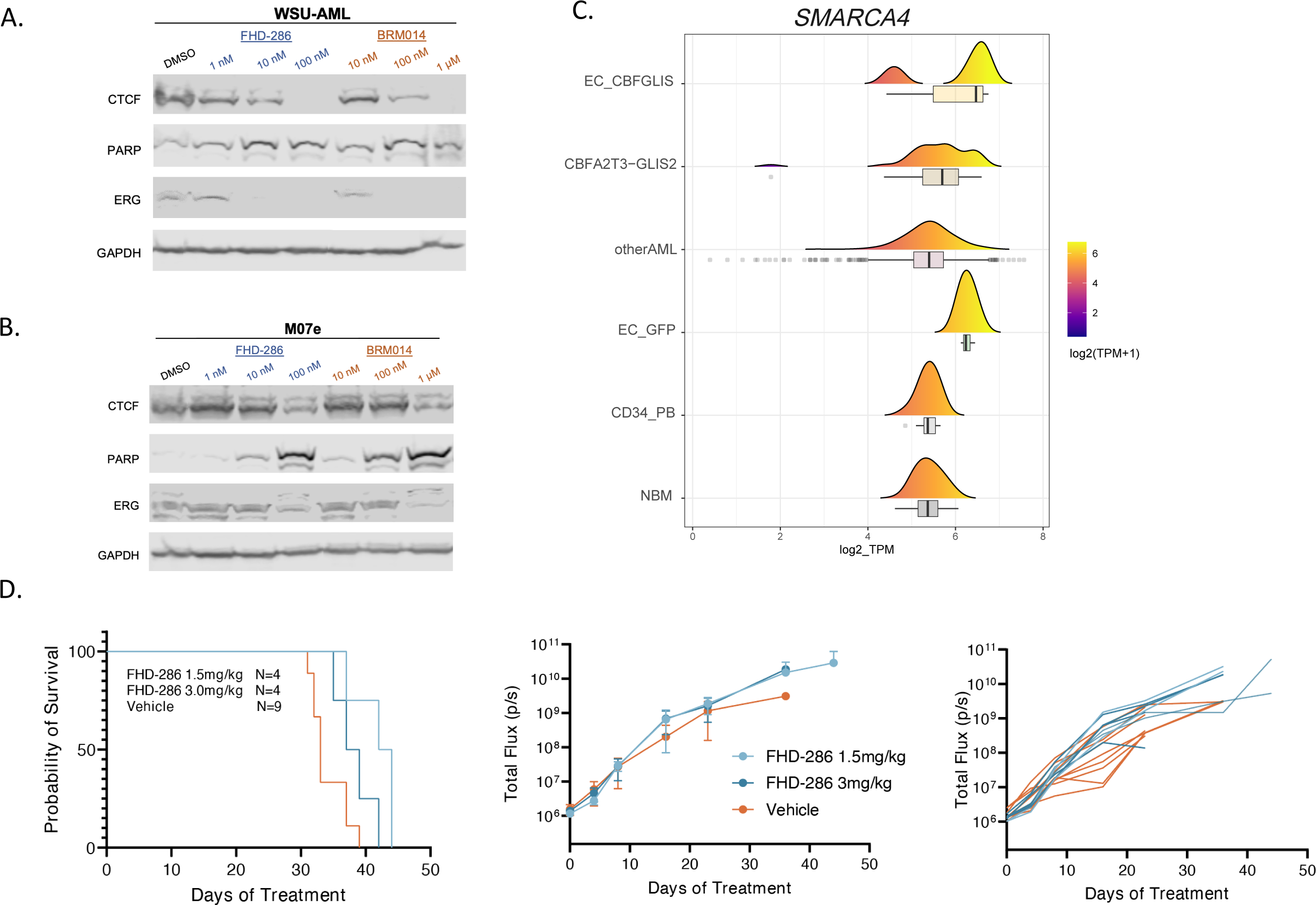
**A.** Western blotting for indicated factors in C/G AML cell lines treated with BRG1/BRM inhibitors for 48hrs. **B.** Density plot of *SMARCA4* in bulk RNA-seq datasets. **C.** Subgroup survival curves and **D.** group and individual luminescence in WSU-AML xenograft trial.

